# Regulatory perturbations of ribosome allocation reshape the growth proteome with a trade-off in adaptation capacity

**DOI:** 10.1101/2021.08.01.454633

**Authors:** David Hidalgo, César A. Martínez-Ortiz, Bernhard O. Palsson, José I. Jiménez, José Utrilla

## Abstract

Bacteria regulate their cellular resource allocation to enable fast growth-adaptation to a variety of environmental niches. We studied the ribosomal allocation, growth and expression profiles of two sets of fast-growing mutants of *Escherichia coli* K-12 MG1655 in glucose minimal medium. Mutants with only 3 of the seven copies of ribosomal RNA operons grew faster than the wild-type strain in minimal media and show similar phenotype to previously studied *rpoB* mutants. Higher growth rates due to increased ribosome content affected resource allocation. Expression profiles of fast-growing mutants shared downregulation of hedging functions and upregulated growth functions. Mutants showed longer diauxic shifts and reduced activity of gluconeogenic promoters during glucose-acetate shifts, suggesting reduced availability of the RNA Polymerase for expressing hedging proteome. These results show that the regulation of ribosomal allocation underlies the growth/hedging phenotypes obtained from laboratory evolution experiments. We show how two different regulatory perturbations (rRNA promoters or *rpoB* mutations) reshape the proteome for growth with a concomitant fitness cost

**Highlights:** Mutants with only 3 ribosomal operons grow faster than wild-type in minimal medium

Δ4 *rrn* and *rpoB* mutants share phenotypic traits

Faster growth of mutants is achieved by increased ribosome content

Fast-growing mutants display reduced hedging expression and adaptation trade-offs

Despite similar ribosomal content in rich medium the mutants present growth defects

## Introduction

Cellular growth is the result of a highly regulated process that demands the allocation of cellular resources towards different tasks. Bacteria have evolved to regulate resource allocation to maximize survival under harsh and changing environments. During fast exponential growth ribosomal RNA represents ~86% of the total RNA, ribosomes constitute over 30% of the biomass (Dennis and Bremer, 2008), and translation uses ~40% of the cellular energy (Wilson and Nierhaus, 2007). Therefore, the regulation of ribosomal content is central to bacterial physiology.

The main control step in ribosome biogenesis is the transcription of ribosomal RNA (rRNA) (Dennis et al., 2004), which hoards much of the available RNA polymerase (RNAP). RNAP is central to the regulation of the ribosomal allocation, it is the target of the (p)ppGpp stringent response and fast-growing strains selected by Adaptive Laboratory Evolution (ALE) often present mutations in the *rpoB* or *rpoC* genes that code for the core RNAP (Phaneuf et al., 2019). ALE selects for RNAP mutants that grow faster than the parental strain in minimal media, display adaptation trade-offs (such as the glucose-acetate diauxic shift) and slower growth in rich media. The effects of these mutations have been attributed to the reprogramming of the whole transcriptome, and the reallocation of resources from downregulated hedging functions to growth, however their effects on the regulation of the ribosome allocation have not been studied (Cheng et al., 2014; Conrad et al., 2010; Utrilla et al., 2016).

A clear correlation between copy number of ribosomal RNA operons, growth rate and growth efficiency has been established for many bacterial species (Roller et al., 2016). In the model bacterium *Escherichia coli* rRNA is transcribed from seven ribosomal operons (*rrn*A, B, C, D, E, G and H) that provide the necessary amount of ribosomal transcription for fast growth. However, it has been shown through progressive deletions that *E. coli* can survive with only one of the seven (Asai et al., 1999a). The resulting ribosomal operon elimination mutants have been studied in rich medium, showing reduced growth rate, increased adaptation time in glucose-LB shift-up assays and a modest reduction in RNA content (Asai et al., 1999b; Condon et al., 1995). Previous studies in *E. coli* have demonstrated that not all the ribosomal operons have the same transcription strength in different conditions, even though the structure of the *rrn* promoters is similar, the genomic location and slight differences in the promoter sequences result in different promoter strength and sensitivity to stringent response (Kolmsee et al., 2011; Maeda et al., 2015). Also, there is evidence showing that poor nutrient conditions produce heterogeneous populations of ribosomes that regulate the expression of stress response genes (Kurylo et al., 2018).

It has been recently shown that the cost of the translationally inactive (unused) ribosomes, present in *E. coli*, may reduce the growth capacity in a static environment but can be useful in a dynamic or changing environment (Dai et al., 2016, Mori et al., 2017). A negative correlation between growth rate and adaptation rate has been recently observed (Basan et al., 2020). This evidence suggests that a fraction of inactive ribosomes may confer fitness advantages in changing conditions, such as those found in nature, however, under constant environments the inactive ribosomes may constitute a burden that constrains the maximum achievable growth rate. An open question regarding the mechanism of growth rate increase of ALE selected mutants in constant minimal media conditions, is if they reduce the amount of inactive ribosomes. The reduction of costly inactive ribosomes could increase growth rate in constant environments but would result in a loss of capacity to resume growth under changing conditions.

Here we studied the allocation of ribosomal resources in two sets of strains with regulatory perturbations that increase growth rates in minimal media. Two mutants with the three stronger ribosomal operon (SQ53: *rrn*CEH, SQ78: *rrn*BCH) expression resulted in higher resource allocation to ribosome synthesis for growth and decreased adaptation capacity. These results are similar to the optimized growth strategies displayed by fast-growing RNAP mutants obtained by ALE (*rpoB*E546V and *rpoB*E672K), which highlights common and specific mechanisms of regulatory reprogramming of ribosomal allocation.

## Results

### The effect of *rrn* operon number on the growth phenotype in different environments

Mutants with deletions of ribosomal RNA operons have been constructed previously (Condon 1993, Asai et al., 1999, Goyfry et al., 2015, Quan et al., 2015, Levin et al., 2017). In general, such studies focused on broad physiological parameters in rich medium, rRNA gene exchange between species and rRNA gene copy implications in antibiotic resistance. Nutrient rich media promote minimal levels of stringent response (through ppGpp, Irving et al., 2021), a global gene expression regulation primarily targeting transcription. The implications of the reduction of ribosomal RNA operons in the general cellular physiology and molecular response in minimal media, where ppGpp levels can be four times higher than in rich medium (Dennis and Bremer, 2008, Zhu et al., 2019) are poorly understood. More so from the perspective of analyzing the regulation of growth phenotypes of the mutants across different environments. To this end we grew the Wild Type (WT: *E. coli* MG1655) strain and the mutants lacking one to six *rrn* operon copies (Table 1, Quan et al., 2015) in rich and minimal media supplemented with different carbon sources. The results showed that while there is a positive correlation between growth rate and number of *rrn* operons in rich medium (LB with glucose and M9 with glucose supplemented with casamino acids, CAA) (Figure 1A), in minimal media with different carbon sources there is an optimal amount of *rrn* operons that maximizes the growth rate (3 *rrn* operons; SQ53: *rrn*CEH, SQ78: *rrn*BCH) and this number is different from that of the WT strain (7 *rrn*, Figure 1B, Supplementary Table 1).

**Table 1.** *rrn* deletion strains. Strain genotypes. Mutants derived from the K-12 MG1655 strain contain deletions of one or several *rrn* operons. Each *rrn* operon contains a copy of the *rrn*, *rrl* and *rrf* genes. In the intergenic and distal positions of these operons there are tRNA genes for different amino acids. Strains may be complemented with a plasmid containing tRNA genes when needed (ptRNA67, SpcR).

| Strain | Genotype | *rrn* in chromosome |
| --- | --- | --- |
| *E. coli*<br>MG1655 | *ilvG rfb-50 rph-1* | 7: ABCDEGH |
| SQ37 | $\Delta rrnE$ | 6: ABCDGH |
| SQ40 | $\Delta rrnEG$ | 5: ABCDH |
| SQ49 | $\Delta rrnGBA$ | 4: CDEH |
| SQ53 | $\Delta rrnGBAD$ | 3: CEH |
| SQ78 | $\Delta rrnGADE$ | 3: BCH |
| SQ88 | $\Delta rrnGADEH$(ptRNA67) | 2: BC |
| SQ110 | $\Delta rrnGADBHC$(ptRNA67) | 1: E |

**Figure 1.**
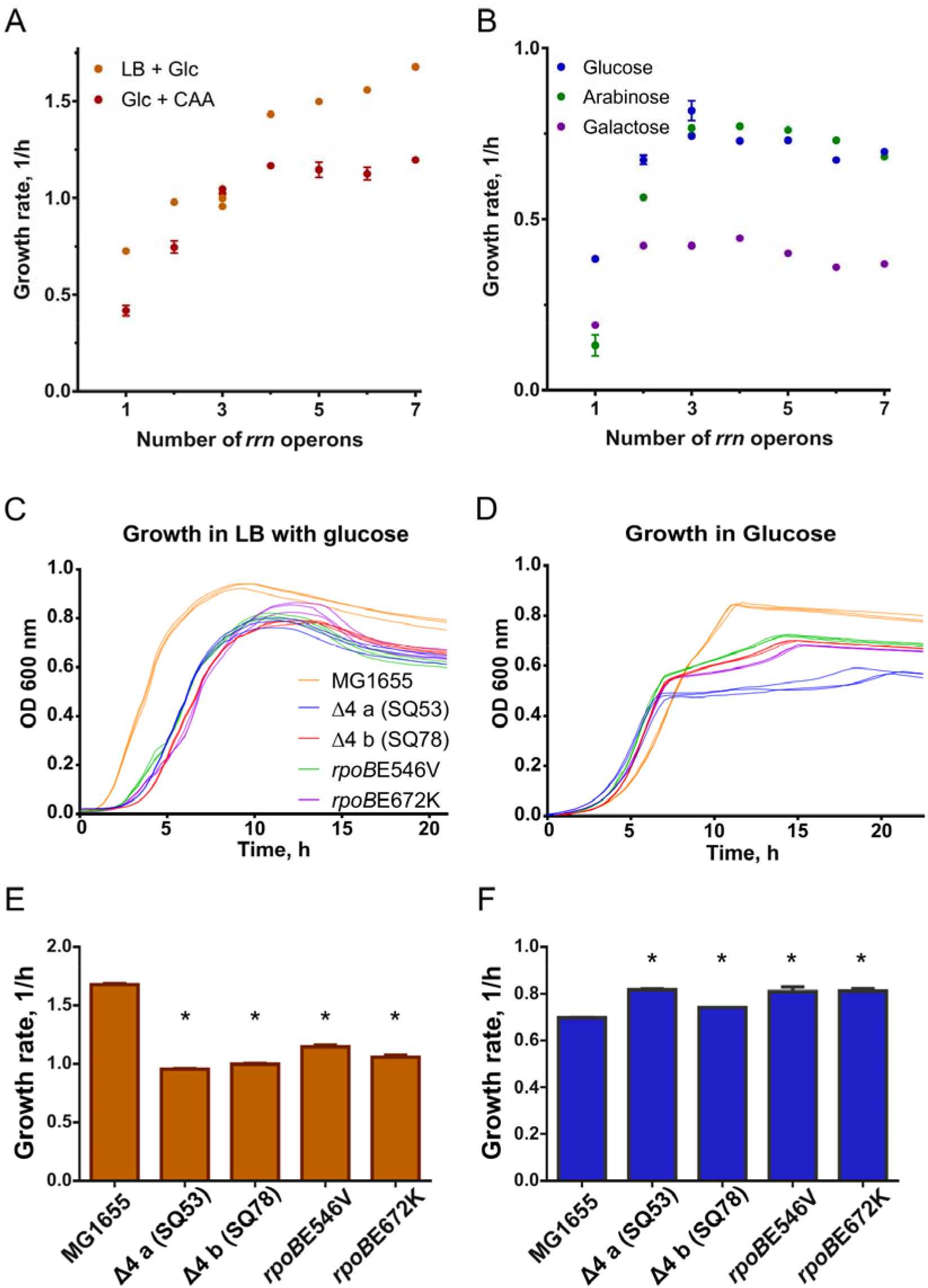
The effect of *rrn* operon number on the growth phenotype in different environments. A, the effect of *rrn* operon copy number on the growth rate in rich media (LB with glucose and glucose supplemented with casamino acids). B, the effect of *rrn* operon copy number on the growth rate in minimal media (glucose, arabinose and galactose). C, Triplicate growth dynamics of selected *rrn* strains and two *rpoB* mutants in LB with glucose. D, Triplicate growth dynamics of selected fast strains in glucose. E, Growth rates of selected strains in LB supplemented with glucose. F, Growth rates of selected strains in glucose minimal medium. Error bars represent standard deviations from nine replicates; some are smaller than the size of the marker. * denotes significant difference with 95% confidence. See supplementary table 1.

Growth dynamics in rich medium and minimal media are shown in Figs. 1C and D, respectively. During our experiments in glucose M9 medium, where acetate is typically produced as an overflow metabolite and latter consumed in a second growth phase, we observed that faster *rrn* mutants showed a diminished ability to resume exponential growth once glucose had been exhausted (Fig. 1D, Fig. S1), while the WT strain transitioned swiftly through the glucose-acetate shift. At this point we recognized similarities between the phenotype exhibited by *rrn* mutants and two previously reported strains (*rpoB*E546V and *rpoB*E672K, Utrilla et al., 2016) each carrying a point mutation in the *rpoB* gene of the RNAP. Both types of mutants grew faster in glucose minimal media, showed glucose-acetate growth shift defects, and grew slower than the WT in rich media, such as glucose + CAA and LB with glucose (Figures 1C, D and E).

### The ribosome production and the growth rate change proportionally to achieve faster growth in minimal media in all mutants

Recently many studies have pointed to the fact that reserve ribosomes and translation efficiency may play a fundamental role in the observed maximum growth rate in a given condition (Dai et al., 2016). Since we studied mutants with a reduced ribosomal operon copy number that grow faster than the WT strain, we hypothesize that a reduced amount of ribosomal content plus a higher translation efficiency may be a possible mechanism for the increased growth rate of the mutants in minimal media. Therefore, one of the main questions we addressed in this work was whether ribosome production in the studied mutants changes in the same magnitude as the WT in different growth environments. To measure ribosome content in the studied strains we used a previously reported *in vivo* system to indirectly monitor ribosome concentration. A ribosomal protein-GFP fusion was used to replace the native *rplI* gene (Fig. 2A), whose protein product is a component of the large subunit of the ribosome (Kim et al., 2021).

**Figure 2.**
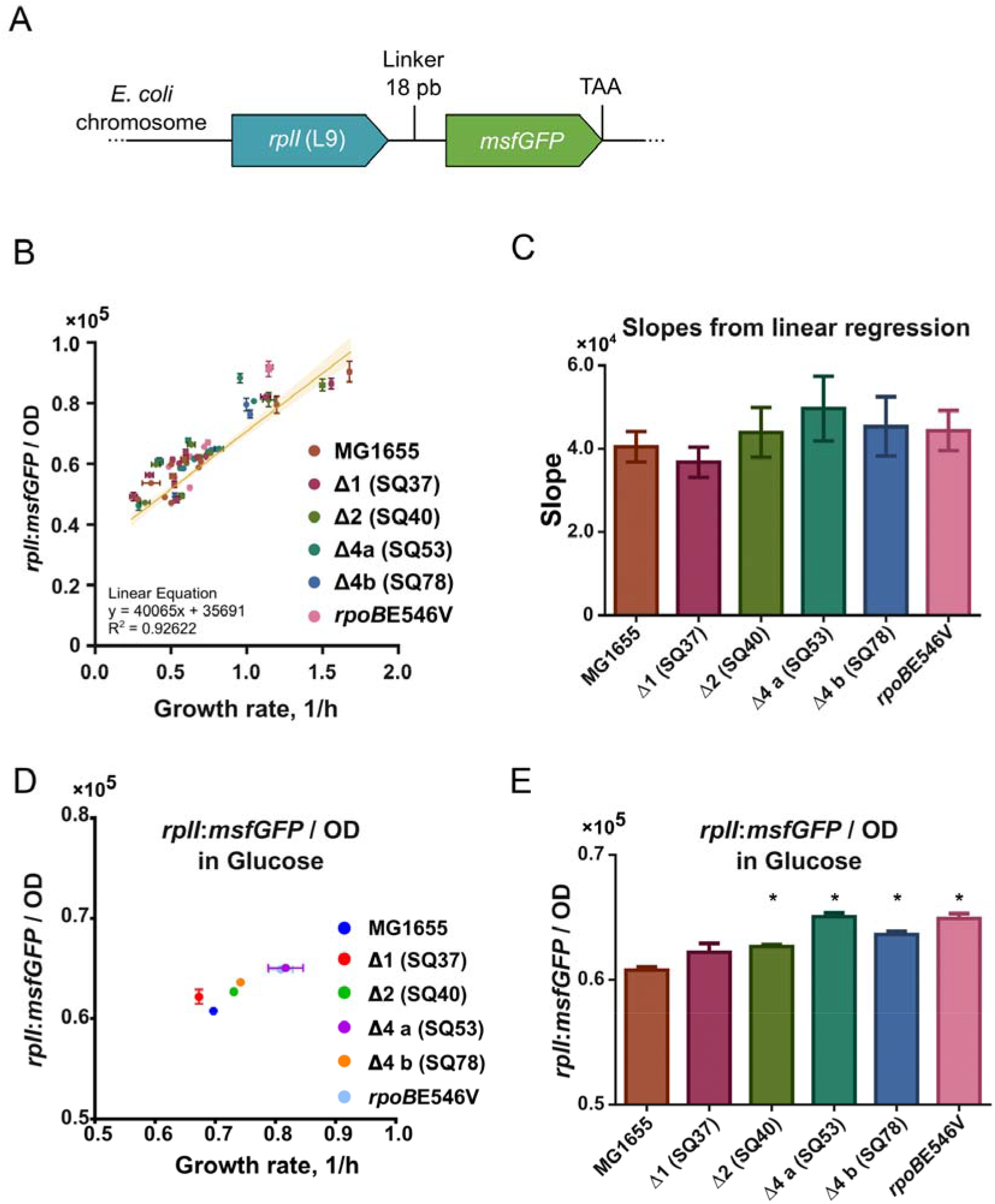
The ribosome production and the growth rate change proportionally in all mutants. A, The ribosomal protein-GFP fusion in the chromosome. An 18 bp linker (coding for amino acids SGGGGS) is placed between the genes (Not to scale). The fused genes encode the ribosomal L9-msfGFP chimeric protein. TAA denotes end of translation. B, Linear regression of the normalized *rplI*:msfGFP fluorescence as a function of growth rate for all strains in several conditions. The yellow line represents the regression for all strains data combined. Each strain’s corresponding points are categorized by color (see legend). Error bars are the standard deviation from triplicates. The regression line’s shadow is the 95% confidence. C, Comparison of each strain’s individual normalized *rplI*:msfGFP slopes obtained by linear regression. No significant differences between any mutant and MG1655 (95% confidence) were found. D, Normalized fluorescence for strains grown in glucose minimal medium as a function of growth rate. E, Normalized fluorescence comparison between mutants and the WT. * denotes significant difference with 95% confidence. See figures S2, S3 and S4.

We measured ribosomal content across 11 media through *rplI:msfGFP* fluorescence. Ribosomal-fluorescent protein fusions have been reported to have a linear correlation with rRNA content (Failmezger et al., 2017). Maximum growth rates (μ) ranged from 0.25 to 1.67 h^-1^. We plotted the normalized *rplI*:*msfGFP* fluorescence during exponential growth as a function of growth rate (Fig 2B); we observed that regardless of the number of rRNA deletions, the slope of the regression of ribosomal content and the and μ is maintained (Fig. 2C). The results also showed that the mutants that grew faster than the WT strain in the same environment did so by increasing the ribosomal content at a similar magnitude than the WT strain for different environments (data points fell on the same linear trend as the WT). Observations in minimal media show that the increase in growth rate derives from an increase in ribosomal content and not from a reduction of inactive ribosomes or increased ribosomal efficiency. For example, in glucose minimal medium an 4.5-6.5 % (rplI:msfGFP/OD) increase in the mutants corresponds to a 6-17% higher growth rate (Fig. 2D and E).

Previous evidence (Kolmsee et al., 2011, Maeda et al., 2015) showed that the promoters of each rRNA operon have different strengths and sensitivity to environmental and regulatory signals such as (p)ppGpp (Fig. S2). Utrilla et al, showed that even though the *rpoB*E546V and E672K mutations are not in the specific (p)ppGpp binding site, they belong to the same “structural community” where molecular motions of residues are strongly correlated. They observed that these mutations perturb the stability of the RNAP open complex in the presence of (p)ppGpp perturbing the response to stringent response with implications in its affinity for sigma factors, ribosomal promoters and other regulatory effects at the transcriptional level. Since both set of mutants increase ribosomal content proportional to the growth increase in 11 evaluated media (Figs S3 and S4 A-H), we postulate that the mechanism for doing so is the stronger expression of ribosomal RNA resulted from two different molecular mechanisms: the use of the stronger set of ribosomal promoters in SQ53 and SQ78 strains and the perturbed stability of the RNAP open complex in *rpoB* mutants.

### Mutants display reduced growth and same ribosomal content than the wild type in rich medium

In this study we investigated the growth increasing strategies due to the perturbation of ribosomal allocation regulation in minimal media. However, our growth characterization shows that in LB + glucose medium the WT growth rate increased by 1.4-fold compared to that in glucose minimal medium, whereas the mutants increased it by only 0.17-0.41 fold. It was previously reported that RNAP mutations in subunits of the RNAP (*rpoB* and *C*) reduce growth rate in rich media (Utrilla et al., 2016, Conrad et al., 2010). To better address ribosomal content in LB + glucose and glucose minimal medium, in addition to normalized rplI:msfGFP data (Fig. S4 I and J), we measured rRNA by capillary electrophoresis (Hardiman et al., 2008) (Fig. S5S). The results show that in the WT strain rRNA increased from 72% to 85% of total RNA when comparing growth in glucose minimal vs glucose rich media. Mutants with higher rRNA in minimal media (74-79%) present slightly less (not significant) amount of rRNA in rich medium than the WT. These results show that in an environment with negligible ppGpp concentration (rich medium), rRNA transcription reaches its homeostatic level, but such hoarding of RNAP may affect global gene expression

### Faster growth phenotypes show a reduced adaptation capacity

In *E. coli* aerobic cultures, growth in glucose minimal medium is accompanied by acetate production (Enjalbert et al., 2015). Two well-defined growth phases can be differentiated in these cultures with a maximum growth rate in glucose and another in acetate (Fig. 3A). We observed that mutants displayed a reduced capacity to adapt and grow in the acetate growth phase (after glucose depletion, Fig. S1). Figure 3B shows the maximum growth rates (μ) before (lilac) and after (green) the diauxic shift. While fast growing mutants (SQ53: *rrn*CEH, SQ78: *rrn*BCH and both in *rpoB*) had a higher growth rate during the glucose phase, they were slower than the WT after the shift (only 28% of the growth rate achieved by the WT for SQ53 and *rpoB*E546V). SQ110, with only one *rrn* copy, grows so poorly that it did not even have a diauxic shift. Regarding the time required to reach the post-shift maximum growth rate, our data showed a delay correlated with the number of rrn deletions (Fig. 3C). However, this may depend on the precise deletions as exemplified by SQ53 which was particularly slow in contrast to SQ78 despite both containing 3 *rrn* copies (see discussion below). Additionally, both *rpoB* mutants showed a similar diauxic behavior to that of SQ78, though with slight differences in the post-shift growth rate which decreased by 28% in *rpoB*E546V and 68% in *rpoB*E672K. These observations show an important phenotypic trait in the fast-growing mutants: they took longer to adapt to a new carbon source and resume exponential growth in acetate and their growth rates were lower than MG1655 in the postshift phase.

**Figure 3.**
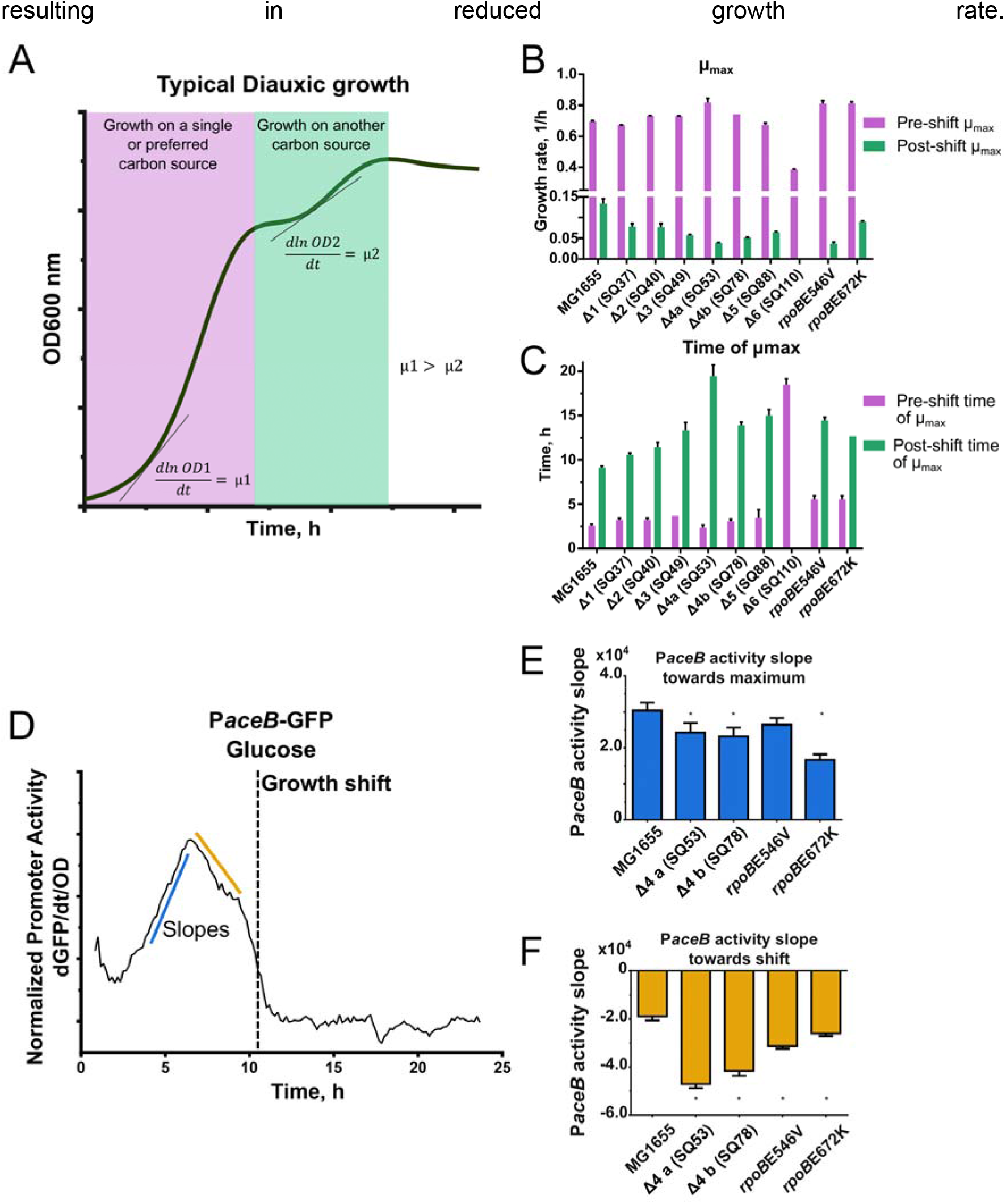
Faster growth phenotypes show a reduced adaptation capacity and the *aceB* promoter activity dynamics. A, Typical bacterial diauxic growth diagram. First, a single carbon source is consumed, reaching a growth rate (μ1). After a pause, another carbon source is consumed and exponential growth is resumed, reaching a second growth rate (μ2) lower than the first one. B, Maximum growth rates (μ) in glucose before (lilac) and after (green) the growth shift. C, Time at maximum growth rates (μ) in glucose before (lilac) and after (green) the growth shift. D, Dynamics of the normalized activity of the *aceB* promoter (dGFP/dt/OD) in MG1655. Dashed line represents the time of growth shift. Red lines represent activity slopes analyzed in panels E and F. E, Comparison of normalized *PaceB* activity slope towards maximum between mutants and MG1655. * denotes significant difference with 95% confidence. F, Comparison of normalized *PaceB* activity slope after maximum between mutants and MG1655. * denotes significant difference with 95% confidence. See also figures S6 and S7.

### *aceB* promoter activity dynamics

The genes *aceA* and *aceB* are essential for growth in acetate. They are responsible for the glyoxylate shunt in the tricarboxylic acid cycle and show a strong transcriptional response in the glucose-acetate shift (Enjalbert et al 2015). In order to test the regulatory response of the *aceBAK* operon in the fast-growing mutants, we used a reporter plasmid containing the *aceB* promoter controlling the expression of GFP (Zaslaver et al., 2006). The normalized Promoter Activity (PA), a measure of RNAP transcription at a certain promoter, was monitored *in vivo* over time in M9-glucose 4 g/L cultures (dGFP/dt/OD, Fig. 3D). In those conditions, two clear phases were observed: a) a gradual increase in *aceB* PA to reach a maximum value of activity hours before the growth shift, and b) a decrease in *aceB* PA towards the transition to acetate growth. The slopes for each phase of this activity dynamics (blue and yellow lines) were determined by a linear regression (Fig. 3E and F). There was a 12-20% reduction in the SQ53 *aceB* PA slope towards its maximum and 54% of the WT PA for the *rpoB* E672K mutant. After this point, the PA slope fell abruptly for the studied mutants with a reduction ranging from 2.47 to 1.37-fold of the WT *aceB* PA. Since the concentration of acetate in each culture may be different and could affect the regulatory response, we also measured the *aceB* PA dynamics in media with glucose-acetate mixtures showing results consistent with those mentioned above: a lower slope of the *aceB* PA during the first phase and a higher negative slope during the second growth phase (Figures S6 and S7). These results showed a reduced transcription of the *aceBAK* operon in both the rRNA and RNAP mutants and suggested a reduced availability of the RNAP to transcribe the genes needed to efficiently adapt to a new carbon source that resulted in a loss of adaptation capacity in the fast-growing mutants.

### The transcriptional profiles of *rrn* and *rpoB* mutants show common differentially expressed functions

After assessing similarities during growth associated with the RNAP in both types of mutants, we compared the transcriptomes in glucose minimal medium of two selected fast strains: SQ53 and the previously reported *rpoB* E672K mutant. Increased growth rate of *rpoB* mutants in glucose minimal medium has been attributed to the reduction of hedging functions, having their associated cellular resources being channeled to growth functions (Kim et al., 2020; Utrilla et al., 2016). Results for SQ53 showed 104 Differentially Expressed Genes (DEGs), of which 74 are downregulated (Figure 4A). Broadly, the downregulated genes belong to hedging functions such as oxidative and osmotic stress, acid resistance, general stress responses and a portion of genes of unknown function (Figure 4B). For the 30 upregulated genes we observed the majority of growth functions such as carbohydrate transport and metabolism, amino acid uptake, and metabolism and protein synthesis. When we compared the expression profile changes among SQ53 and *rpoB* mutants we found similarities in terms of the broad categories of DEGs, such as osmotic and oxidative stress, acid resistance, ion transport and homeostasis and genes with unknown function.

**Figure 4.**
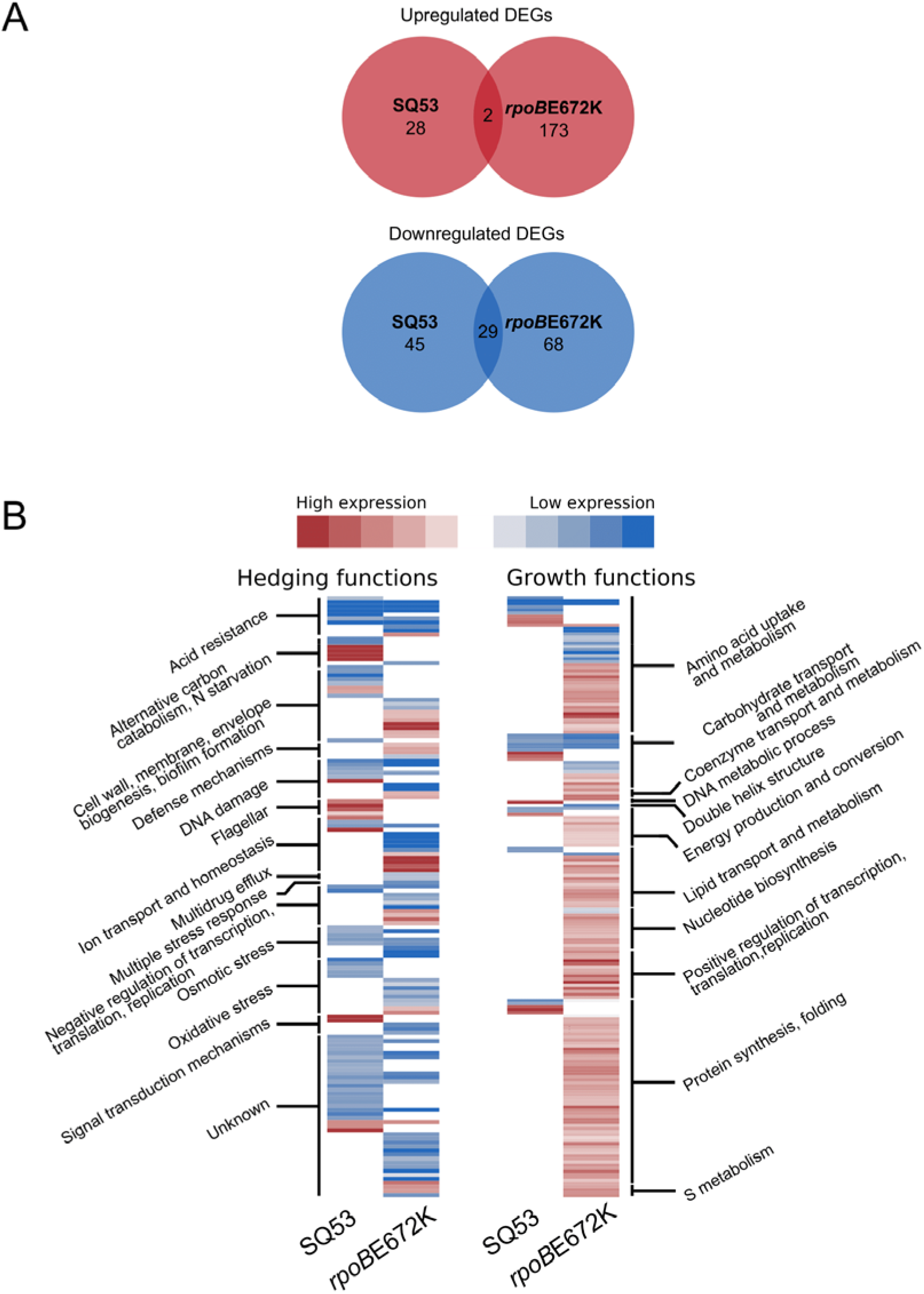
The transcriptional profiles of *rrn* and *rpoB* mutants show common differentially expressed functions. A, Number of upregulated (red) and downregulated (blue) genes for SQ53 and *rpoB*. Intersections denote the number of shared genes between mutants. B, Heatmap showing expression levels of hedging and growth functions for SQ53 and rpoB672. Genes were categorized by the functions shown. See supplementary file.

We further analyzed the data by performing a proteome balance calculation to estimate the contribution in mass by each DEG (Lastiri-Pancardo et al., 2020, see methods). While a given gene may show differences in its transcription, it is the size and copy number of the corresponding protein that has the largest impact on the cellular resource and energy budget (Lynch and Marinov, 2015). From these calculations we observed that most of the impact on the cellular budget was associated with a few proteins showing a Pareto-shape response regardless of the mutant. This means that 80% of the change in cellular budget associated with gene expression comes from just a few genes (Figure 5). For upregulated genes, most categories corresponded to amino acid uptake, protein synthesis, carbohydrate transport and metabolism and ion homeostasis. For downregulated genes, the associated categories are more heterogeneous. Within the genes with more impact, functional categories corresponded to oxidative and osmotic stress, DNA damage and metabolic process, ion homeostasis and defense mechanisms. Another important observation derived from these calculations is that the estimated proteome change was higher for upregulated functions compared to the down regulated ones. Upregulated genes for SQ53 contributed three times the mass of those downregulated (6.27 vs 2.09 fg., respectively). For the *rpoB*E672K mutant the difference is more prominent, as upregulated genes contributed 5.97 times more than downregulated genes (52.43 vs 8.76 fg., respectively). This means that the estimated increase in the growth proteome for both studied mutants is higher than the reduction of the hedging functions. This may explain the better capacity to grow in minimal media but the reduced ability to shift conditions. The mutants show a larger and more specialized proteome that prevents them from being prepared for environmental adaptations.

**Figure 5.**
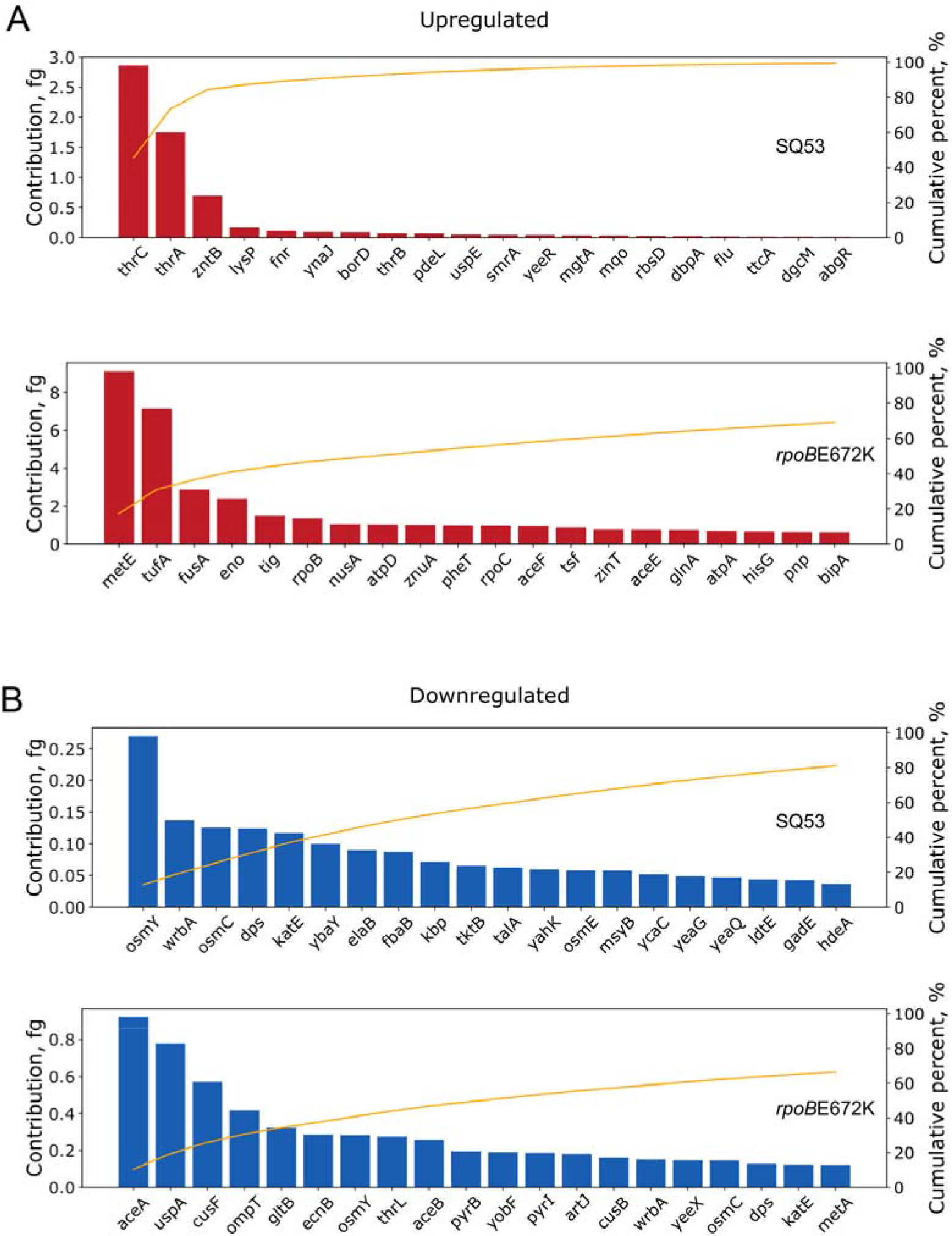
Proteome mass contribution in femtograms for DEGs. A, B Proteome mass contribution in femtograms for up and downregulated genes for each strain, respectively. The yellow line represents the cumulative percent (right axis) for the 20 genes with the largest contribution.

### Greed versus fear response and the proteomic budget of fast-growing mutants

In order to get a deeper understanding of the transcriptomic changes in fast growing strains we used the iModulon classification of the DEG data sets. iModulons are a set of genes obtained by independent component analysis of transcriptomic data sets that reflect the transcriptional regulatory network of *E. coli* (Sastry et al., 2019). We found significant and conserved changes in few iModulons: there was a general increase of the iModulons related to growth (pyruvate, BCAA-2), and a general reduction to the “fear” iModulons (*rpoS*) (Figs. 6A and S8). Tan et al 2020 reported the expression patterns in different heterologous protein production strains and observed that those proteins that elicited a weaker expression of fear iModulon were more productive for heterologous protein production. We therefore assessed the availability of cellular resources for recombinant gene expression using the isocost lines methodology that measures the protein budget on the basis of the relationship of two coexpressed fluorescent reporters (Gyorgy et al., 2015). Even though the fast-growing strains show a higher expression of the translation machinery iModulons (greed) and reduced the expression of the *rpoS* iModulons (fear), we found a reduced proteomic budget in these strains (Figure 6B). Recently, it has been observed that in a defined environment fast growth and unnecessary protein production are inversely correlated (Kim et al., 2021; Scott et al., 2010). Therefore, the increased allocation of resources to growth obtained in these strains is not correlated with a higher budget for a synthetic function. Taken all together, our results suggest that the similar phenotype of the RNAP and rRNA mutants is the result of different expression profiles with similar categories of genes being differentially expressed.

**Figure 6.**
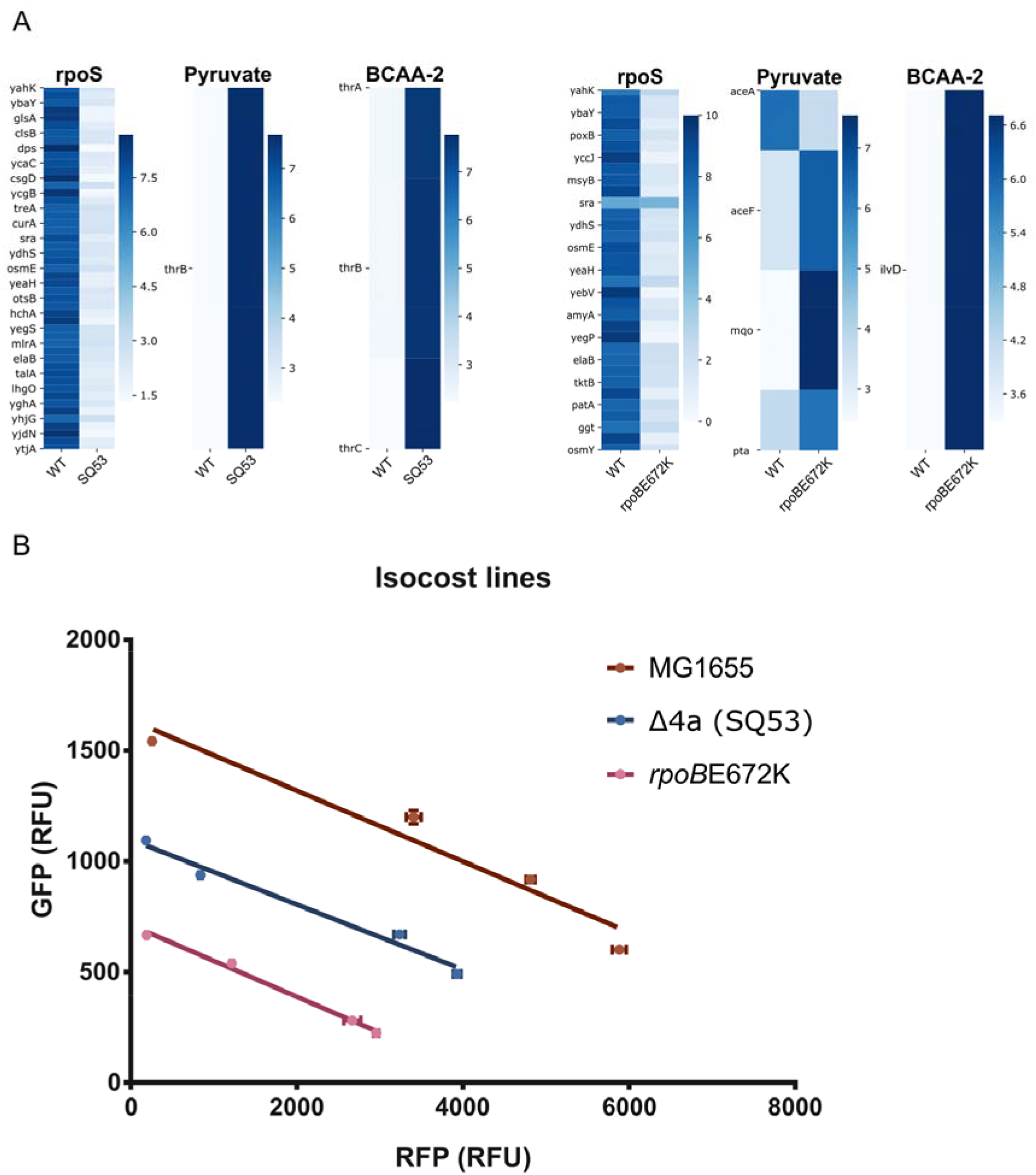
Greed versus fear response and the proteomic budget of fast-growing mutants. A, DEGs categorized by iModulons for SQ53 and *rpoB*E672K. Color bar indicates fold change expression level. B, Isocost lines indicating differences in cellular budget between mutants and the WT. See Fig. S8.

## Discussion

The bacterial growth rate is the result of a complex regulatory interplay that fine tunes the allocation of cellular resources such as the ribosomes. Several works have proven that the growth rate in a determined environment can be increased if the proteome is optimized, by regulatory and other mutations, to that specific environment (Cheng et al., 2014; Heckmann et al., 2020; LaCroix et al., 2015). Indeed, rewiring *E. coli* transcription networks with fusions of promoters and master regulators (e.g. *arcA, crp, rpoD*) upregulated ribosomal-associated genes and increased growth rates (Baumstark et al., 2015). Here, we show that different regulatory perturbations that increase ribosomal synthesis result in higher growth rates in specific environments are also associated with phenotypic and molecular trade-offs.

In this work, we studied *E. coli* mutants lacking one to six copies of the seven ribosomal operons. We found that, while in rich media ribosomal deletion results in a reduced growth, there is a combination of three ribosomal operons that results in higher growth in minimal media. We compared the fast-growing ribosomal mutants to the ALE selected RNAP mutants (*rpoB*E546V and *rpoB*E672K) due to their striking growth phenotype similarity: these strains exhibit increased growth rate in minimal media with longer diauxic shifts and decreased growth rate in rich media. Our data suggests this arises from a regulatory effect on RNAP distribution, being able to increase the expression of rRNA in very specific conditions with faster growth and a reprogramming of the whole transcriptome. Even though some studies point to an increase in ribosomal efficiency or the reduction of reserve ribosomes as a possible mechanism of increase in growth rate (Dai et al., 2016), here we show that the increase in growth rate in the studied mutants was due to the increase in ribosomal content. We attribute this to two different mechanisms with similar consequences: the previously shown differential regulation and strengths of ribosomal operons (Kolmsee et al., 2011; Maeda et al., 2015) or mutations in RNAP that may perturb the sensibility to the (p)ppGpp stringent response. Both mechanisms can perturb the (p)ppGpp - ribosomal allocation regulatory circuit. Therefore, in wild type *E. coli* the (p)ppGpp - ribosomal allocation regulatory circuit is fine tuned to constrain the maximum growth rate with a better adaptation response.

In the specific case of rich medium (LB + glucose) both types of mutants have similar ribosome content (rplI:GFP, %rRNA) as the WT but grow slower (57-68% of the WT growth rate). The low ppGpp concentration (four times less than in glucose minimal media) may drive the increased transcription of rRNA in the remaining rrn operons as has been shown before (except for a 6 rrn deletion mutant, Asai et al., 1999b). To sustain such a level of transcription, various steps in the synthesis of rRNA should occur. Namely, increased elongation rate (ER) and increased elongation initiation at the remaining rrn copies (Zhu et al., 2021). A study of the kinetics of ALE identified *rpoC* mutants of the RNAP shows increased elongation rate in *rrn* transcription units (*rrn*B, Conrad et al., 2010) and a short lived RNAP open complex. Since protein synthesis requires more than just ribosomes (ternary complexes, composed of GTP, charged tRNAs and EF-TU) it is likely that the tradeoff of homeostatic rRNA transcription is a deficit in the expression of necessary genes. The RNAP redistribution due to the adaptive mutation for growth optimization in minimal medium does not imply that this beneficial effect should be reflected also in rich medium. Indeed, Utrilla 2016 reports a modest but significant decrease in growth-associated genes in rich medium. Additionally, RNAP hoarding and perturbation of the elongation rate likely disrupt the coupling of transcription-translation, further affecting gene expression.

Regarding adaptation, we found that mutants that showed faster growth rate in glucose also showed the slowest growth rate in the acetate consumption phase. This agrees with a study (Basan et al., 2020) where cultures with higher pre-shift growth rates show the longest times to resume exponential growth in shift-down experiments due to reduced metabolite pools and enzyme distribution. Furthermore, the *aceB* PA results show that these fast-growing strains allocate less resources to anticipate the carbon source shift. This is very likely due to the sensitivity of the *rrn* promoters to environmental and regulatory signals, the SQ53 (*rrn*CEH) and the SQ78 (*rrn*BCH) strains both harbor the *rrnH* ribosomal operon that has been shown to be upregulated in minimal media (Kurylo et al., 2018). Furthermore, as it has been previously shown, the P1 promoters of *rrn* operons have heterogeneous strength and sensitivity to (p)ppGpp (Kolmsee et al., 2011, Maeda et al., 2015, Fig.S2). The SQ53 mutant has the strongest and least (p)ppGpp sensitive P1 promoter (*rrn*E). The SQ78 mutant, differing only in having *rrn*B instead of *rrn*E, has a more responsive P1 promoter to reduce rRNA transcription. Our results suggest that the sensitivity of the P1 promoters to regulatory signals allows the RNAP to be allocated to the transcription of genes required for shifting growth to a gluconeogenic carbon source in a more timely fashion. In summary, the WT strain with higher *aceB* PA had a shorter time to resume exponential growth compared to the fast-growing mutants with lower *aceB* PA. The fast-growing *rpoB* mutants showed a similar phenotype, and while the detailed molecular mechanism of the regulatory reprograming in these mutants needs to be further studied, overall these results point to a reduced sensitivity to (p)ppGpp that drives RNAP to highly express ribosomal operons with a lower capacity to shift to alternative promoters.

The transcriptome of the fastest growing strain, SQ53, showed a general profile similar to what was previously observed for the *rpoB*E672K strain. There was a conserved response of increased growth functions and a reduction of hedging functions, however the specific genes differentially expressed in the strains were not the same (Utrilla et al. 2016). In this study, we conducted a calculation of the proteomic contribution of each DEG in both mutants compared to the WT strain. We observed that there is a greater increase in proteome related to growth functions than the estimated proteome reduction from downregulation of hedging functions. These results provide new insights into the molecular phenotype of the fast-growing mutants: they produce a larger and more specialized proteome resulting in a tradeoff in adaptation capacity. Here we show that the fast growth phenotype may be associated to a more efficient recruiting of the RNA polymerase to the *rrn* promoters with a concomitant reduced expression of bet-hedging genes such as those of metabolic adaptation to gluconeogenic carbon sources, thus highlighting its evolutionary role for selecting fast growth phenotypes.

Analyzing the transcriptomic changes with the iModulon methodology we identified a general shift in the greed versus fear response; however, we did not observe a larger proteomic budget for a synthetic circuit expression as measured by isocost lines. Even though previous works show that less fear iModulon expression, when expressing unnecessary proteins, results in higher proteomic budget (Ceroni et al., 2015; Tan et al., 2020), our results show the growth rate related greed versus fear balance do not result in increased proteomic budget. Recent evidence shows that a faster growth rate in a given environment results in a reduced budget for a genetic circuit (Kim et al., 2021). From a resource allocation and a cellular minimization point of view (Hidalgo and Utrilla, 2019), the detailed study of these strains is an important effort to understand and harness design principles for tailored cellular chassis construction.

In this work, we took a systems-based approach to study perturbations on rRNA synthesis at a molecular level. We show that increasing ribosomal content resulting from regulatory perturbations increases growth rate in minimal media but has an adaptation capacity cost and less resource availability for engineered functions. Our findings show that the *E. coli* proteome can naturally achieve faster growth rates in minimal media, but its regulatory program limits the ribosomal allocation to channel resources for hedging functions. The understanding of the correlation between growth, gene expression machinery, and its regulation, are of great interest for those studying the general physiology of bacteria and can help to guide successful bioengineering efforts.

### Limitations of the study

Our work directly provides critical and systems-level insights of the growth profile and dynamics including medium-transitions with sensible *in vivo* molecular probes for fast mutants, however, a dynamic measurement of the whole proteome at different points of the diauxic shift could provide insights of the regulatory response at a finer time scale. Regarding the adaptation trade-off, assays evaluating RNAP activity or localization would support reduced RNAP availability as the major contributor for the downregulation of hedging-function genes.

In the case of rich medium, a detailed description explaining the differences between growth rates but similar ribosome content in the mutants compared to the WT would require additional work beyond the scope of this study.

Lastly, this study identifies the molecular factors contributing to faster growths and their adaptation trade-offs. A detailed molecular mechanism showing how two contrastingly different mutations lead to similar phenotypes is yet to be elucidated. Direct ppGpp quantification, which is not trivial, would help to explain such a mechanism.

## Acknowledgements

We thank H. King for rRNA quantification equipment support. We acknowledge the funding provided by UNAM–DGAPA-PAPIIT project IN213420 and Newton advanced Fellowship Project NA160328. D.H. is a doctoral student from Programa de Doctorado en Ciencias Biomédicas, Universidad Nacional Autónoma de México (UNAM), and received a fellowship 508351 from CONACYT. C.M. acknowledges the Programa de Maestría y Doctorado en Ciencias Bioquímicas, UNAM and the masters scholarship 825745 from CONACyT.

## Author contributions

Experiments, data processing, statistical analyses and figures were conducted by D.H. RNA extraction, library construction and initial RNAseq data processing and analysis was carried out by C.M. Critical analysis of experimental data and RNAseq categorization was done by D.H and J.U. *In vivo* ribosomal measurements analysis and isocost lines were carried by D.H. and J.J. The project was conceived by D.H. and J.U. Supervision and guidance by J.U. The manuscript was written by D.H., J.U., J.J. and B.P.

## Accession numbers

The accession number for the RNAseq data of SQ53 and processed files of SQ53 and rpoBE672K reported here is GEO:GSE180830. The accession number for the RNAseq data of rpoBE672K reported in Utrilla et al., 2016 is GEO:GSE59377.

## Methods

### Strains

The WT strain used in this work was *E. coli* K12 MG1655.

The *rrn* deletion strains are derived from *E. coli* K12 MG1655 and were generated by Quan et al. 2015. and obtained through the Coli Genetic Stock Center (CGSC, http://cgsc2.biology.yale.edu/)

Table 1 summarizes the strain’s genotype.

The *rpoB* mutants *rpoB*E546V and *rpoB*E672K were obtained from Utrilla et al., 2016.

### Growth media

Rich complex media used in this work were LB (Sigma), LB with 1 g/L glucose and M9 minimal media with 0.2%casamino acids (cAA) (Sigma) and 4 g/L glucose. The minimal medium used was M9 (33.7 mM, Na2HPO4; 22.0 mM, KH2PO4; 8.55 mM, NaCl; 9.35 mM, NH4Cl; 1 mM, MgSO4; 0.3 mM, CaCl2) supplemented with trace nutrients (13.4 μM, EDTA; 3.1 μM, FeCl3 - 6H2O; 0.62 μM, ZnCl2; 76 nM, CuCl2 - 2H2O; 42 nM, CoCl2 - 2H2O; 162 nM, H3BO3; 8.1 nM, MnCl2 - 4H2O).

All carbon sources (glucose, glycerol, arabinose, galactose, sodium gluconate, xylose, fructose) were at 4 g/L except for acetate and succinate (2 g/L) unless otherwise stated (diauxic shift experiments). Antibiotic concentrations were Gentamicin 25 μg/mL, Spectinomycin (Spc) 50 μg/mL and Kanamycin 50 μg/mL unless otherwise stated.

### Cell growth

Frozen strains in glycerol at −80°C were grown in solid medium and incubated overnight to 24h at optimal temperature. A colony was put into liquid medium with the corresponding antibiotics were needed and incubated overnight to 24h in agitation (37°C)

On the day of the experiment, either plates (50 μL) of flasks (70 mL) were inoculated at 0.05 OD 600 nm.

### Growth-associated calculations and determinations

#### Growth rates

After measuring the OD at 600 nm at 20 minutes intervals in an automatic plate reader (ELx808, Biotek), the growth rates for the plate experiments (3 replicates per plate for 3 independent plates) were determined computationally using an algorithm based on gaussian processes (Swain *et al*., 2016) running on Python. From this analysis, the magnitude and time of the maximum growth rate and of the maximum OD were obtained, as well as the time and magnitude of local growth rate maxima. Additionally, for selected growth media, the time and OD at diauxic changes was analyzed manually.

When needed, fluorescence intensity in plate experiments was also measured using an automatic plate reader (H1, Biotek) at 479 nm excitation and 509 nm emission.

### *rplI*:msfGFP-Tagged strains

Strains were electroporated with the cpEMG *rplI*:msfGFP plasmid and recovered in rich medium overnight. 200 μL of the recovery were plated in LB Km (25 μg/mL). Cointegration in green fluorescent colonies was verified by PCR using primers emg rpli Fwd (5’-agggataacagggtaatctgcgctcgctacctgtccctgct-3’) and yifz emg Rev (5’-gcctgcaggtcgactctagagtatttattgcaagatgtcgaat-3’). Km resistance was not removed since it does not interfere with the rplI:msfGFP fusion. Experiments were run in the absence of the antibiotic.

### Glucose quantification

Glucose quantification was performed with a YSI 2700 Select machine directly on filtered supernatants of cultures grown in M9 medium with 4 g/L glucose sampled at selected times. Calibration of the equipment was done with a standard glucose solution. Each run uses a 20 μL sample.

### rRNA quantification

Based on Hardiman et al., 2007 with slight modifications. Briefly, Samples for total RNA extraction taken from agitated flask cultures at early exponential growth (0.2 OD at 600 nm). 2 mL samples were mixed into glass tubes containing 2 volumes of RNAprotect bacteria reagent solution (Qiagen). After mixing and incubating for 5 min at room temperature, 2 mL aliquots were separated into three 2 mL microcentrifuge tubes and centrifuged at 14 000 rpm for 10 minutes at 0°C). The supernatants were removed, and the cell pellets were frozen at −20 °C.

Selected samples were resuspended in 1 mL of protoplasting buffer (15 mM Tris–HCl [pH 8.0], 0.45 M sucrose, and 8 mM EDTA and stored at 4 °C), and 10 μL lysozyme solution (50 g/L hen egg white lysozyme, Sigma) was added. After gently mixing and incubating on ice for 15 min, the protoplasts were collected by centrifugation (10 000 rpm for 5 min at 0 °C). The supernatant was carefully discarded and the protoplasts were resuspended in 0.5 mL of lysing buffer (10 mM Tris–HCl [pH 8.0], 10 mM NaCl, 1 mM sodium citrate, and 1.5% [w/v] SDS stored at 37 °C). Then 15 μL diethyl pyrocarbonate (DEPC) was added. The resulting preparation was incubated for 5 min at 37 °C and subsequently chilled on ice. After mixing with 250 μL of a saturated solution of NaCl and incubating for 10 min on ice, the protein–SDS-DNA precipitate was collected by centrifugation at 14 000 rpm for 20 min at 0°C. Then 500 μL of the supernatant was transferred to a fresh 1.5 mL microcentrifuge tube and mixed with 1 mL of ice-cold 100 % ethanol. The obtained precipitates were incubated at −80 °C for 30 min prior to centrifugation at 14 000 rpm for 25 min at 0 °C. The nucleotide-containing pellet was washed with 1 ml ice-cold 70 % ethanol and centrifuged at 14 000 rpm for 3 min. The ethanol was removed carefully and the resulting RNA extract was resuspended in 10 μL RNase-free water for immediate analysis (RNA 6000 Nano kit, Bioanalyzer instrument and the Bioanalyzer 2100 software, Agilent).

All solutions were prepared with RNase-free water prepared with 0.1% DEPC in dest. water, incubated overnight at 37 °C, and then autoclaved.

### p*PaceB*-GFP

The Km resistant plasmid containing the aceB promoter fused with GFP was constructed by Zaslaver et al., 2006. Cells were transformed by electroporation following a 1-hour recovery and plated on LB with Km. Fluorescent colonies were selected for the plate assays.

### RNA sequencing

SQ53 and MG1655 cells were cultured at 37°C in 25 mL of glucose minimal medium. During early exponential growth, 3 mL were sampled into a tube and 3mL of RNA protect reagent were added and mixed. After a 5-minute incubation, samples were lysed enzymatically (Lysozyme from chicken egg white, Sigma). RNA from samples was purified using a modified version of the Quick-RNA miniprep kit from Zymo research. RNA integrity of purified samples was verified by gel electrophoresis (1% agarose) using TAE buffer. Then, 10 μL of each replicate at a 500 ng/μL concentration was used as the input material to be separated from ribosomal RNA using the Zymo-Seq RiboFree Total RNA Library kit from Zymo research. The resulting material was used to construct the libraries using the TruSeq Stranded mRNA kit from Illumina and sequenced using the Illumina Next-Seq 500 platform. Libraries were sequenced to 10 million paired reads. These reads were then verified for quality and the adapter sequences were cut. Sequence alignment using the *E. coli* genome (GCF_000005845.2_ASM584v2) as reference was performed using R packages from bioconductor. Tables containing the raw transcripts were used as input for the normalization algorithm and significance tests from the DeSeq2 package, comparing replicates from the WT and mutant strain in the same experiment. Cut values were set to p < 0.1 with a base 2 logarithmic fold change different from 0. Genes passing these criteria were recognized as differentially expressed. The accession number for the RNAseq data reported in this paper is GEO: GSE180830 (https://www.ncbi.nlm.nih.gov/geo/query/acc.cgi?acc=GSE180830).

### Proteome estimation

The proteome estimation calculation, as reported by Lastiri-Pancardo et al. 2020, is based on the number of transcripts (*ri*) and the corresponding protein copy number per cell (*C* cell *i*). It is assumed that the transcripts yield the proteome reported by Schmidt et al. in glucose minimal medium. The translation efficiency rate (*si*) is calculated by

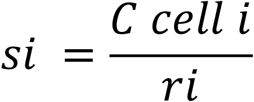

The rate for each protein is then used to estimate the protein copies per cell (*Pi*) for the mutants and transform that to mass with

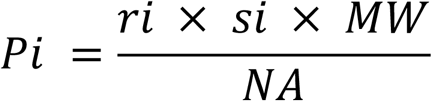

where *Pi* is a given protein *i* contribution in mass, *ri* is the mutant’s transcript abundance in FPKMs, *MW* is the protein’s molecular weight and *NA* is Avogadro’s number (6.022 x 10^23^). For each protein, the difference between mutant and WT Pi is its contribution in mass.

### Gene categorization

Differentially expressed genes from SQ53 and rpoB672 were categorized using available data from ecocyc (https://ecocyc.org/) and uniprot (https://www.uniprot.org/) and compared with previous categorization by Schmidt, 2016. Final categories were manually cured based on the most frequent annotations from available data sources (see supplementary file). Genes with poor or unavailable information regarding function or associated sigma factor were considered in the hedging function group.

## Supplementary material

Supplementary table 1. Related to figure 1. Growth rates, % change and significance. The growth rates from plate assays in rich and minimal media are presented. The % change compared to the WT was calculated and the statistical significance was determined at 95% confidence.

**Figure S1.**
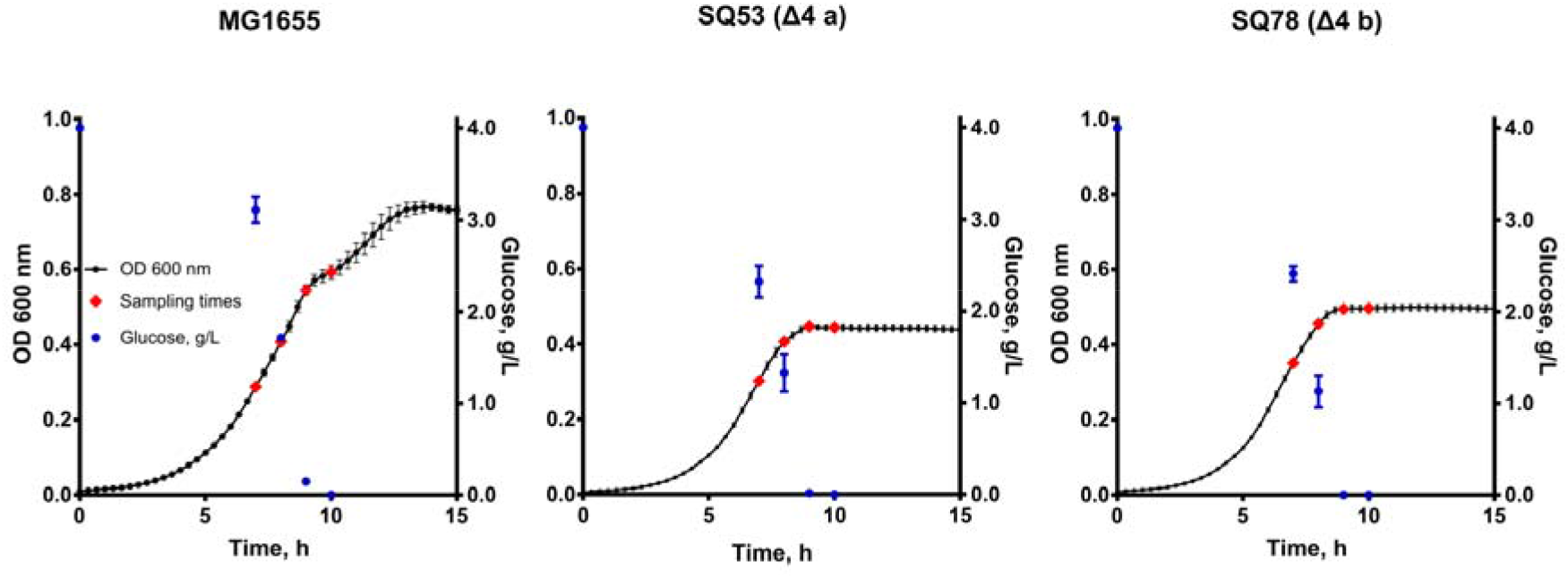
Related to figure 1D and 3. Glucose quantification. Glucose concentration was determined directly in supernatants of samples taken from MG1655, SQ53 and SQ78 during growth on glucose minimal medium. Samples were taken at t = 0h and close to the growth shift. Glucose is depleted at the growth shifts.

**Figure S2.**
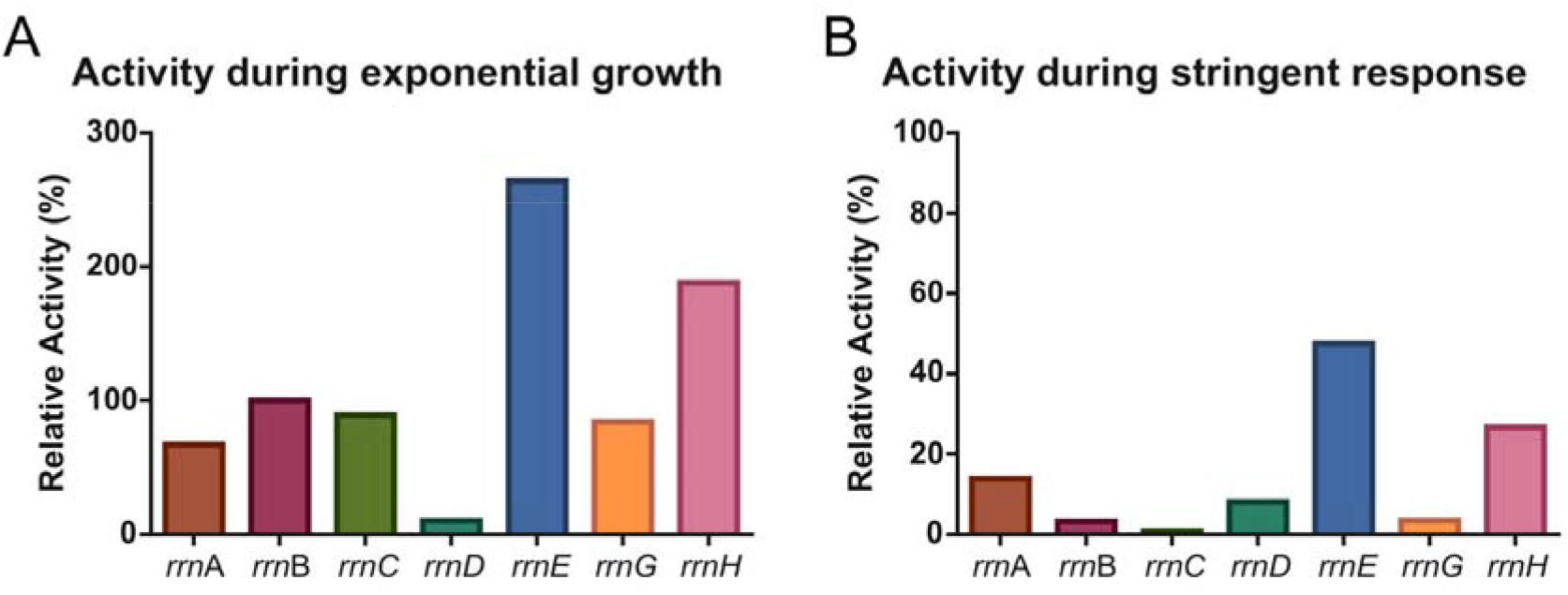
Related to Fig. 1, 2 and 3. Taken and adapted from Kolmsee et al., 2011. Individual strengths of the rrn P1 promoters during A, exponential growth, and B, stringent response, as measured by a stable RNA product and relative to the *rrn*B P1 promoter. Panel A shows the heterogeneous activities among P1 promoters, ranging from 10% (*rrn*D) to 264 % (*rrn*E). Panel B shows reduced P1 promoter activities due to (p)ppGpp action, also elucidating heterogeneous sensitivity to stringent response. SQ53, with 3 rrn copies and having the strongest but also the least sensitive P1 promoter (P1*rrn*E) responds poorly to signals to reduce rRNA transcription (the major effect of stringent response). SQ78, with the same number of rrn copies but differing only in having *rrn*B instead of *rrn*E, has more responsive P1*rrn* promoters to stop rRNA transcription. This allows the RNAP to be allocated in the transcription of other necessary genes for growth in a new carbon source and, hence, has a shorter time to resume exponential growth.

**Figure S3.**
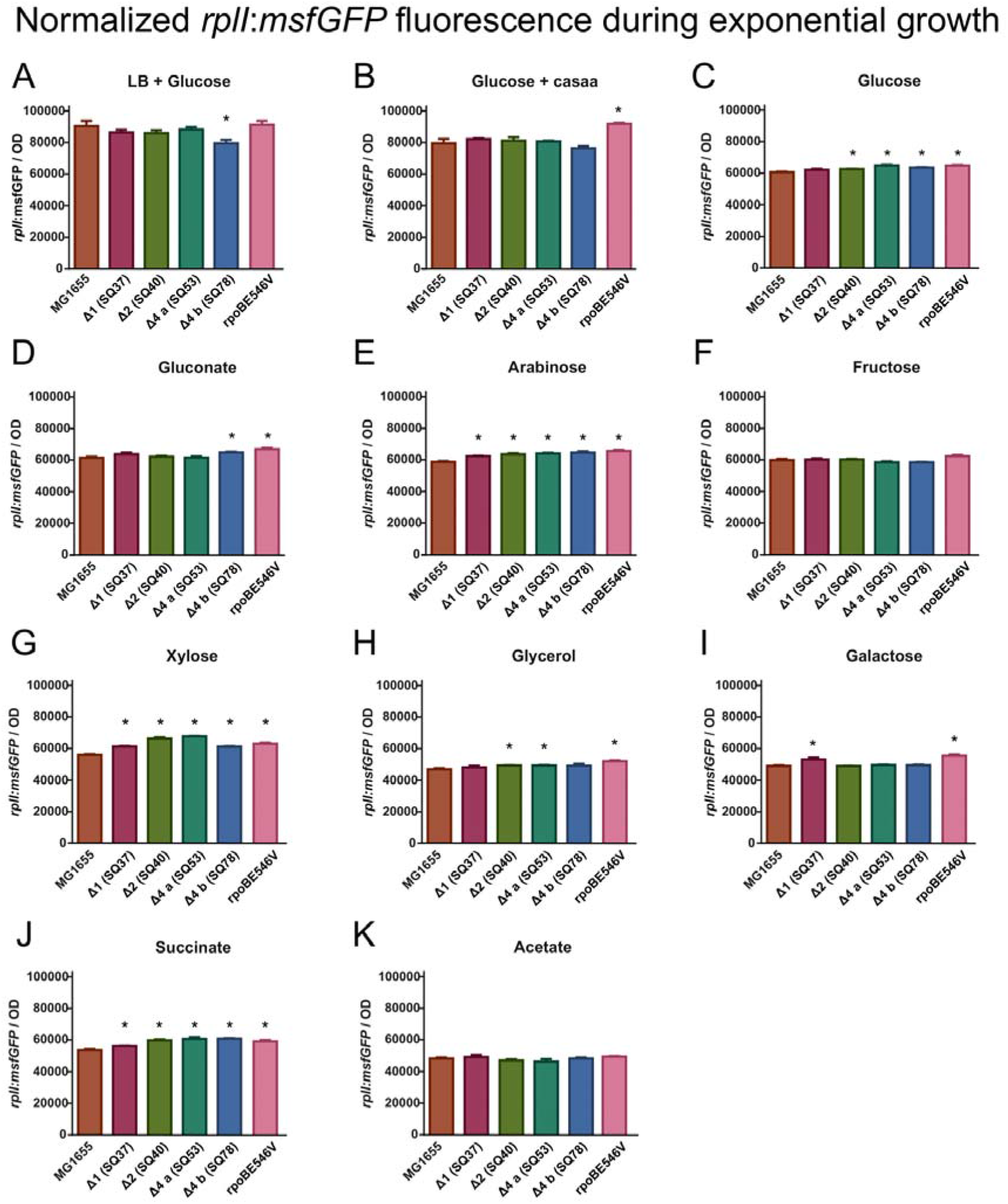
Related to Fig. 2. A-K, Normalized *rplI:msfGFP* fluorescence during exponential growth for mutants grown in different media. * denotes significant differences (95% confidence).

**Figure S4.**
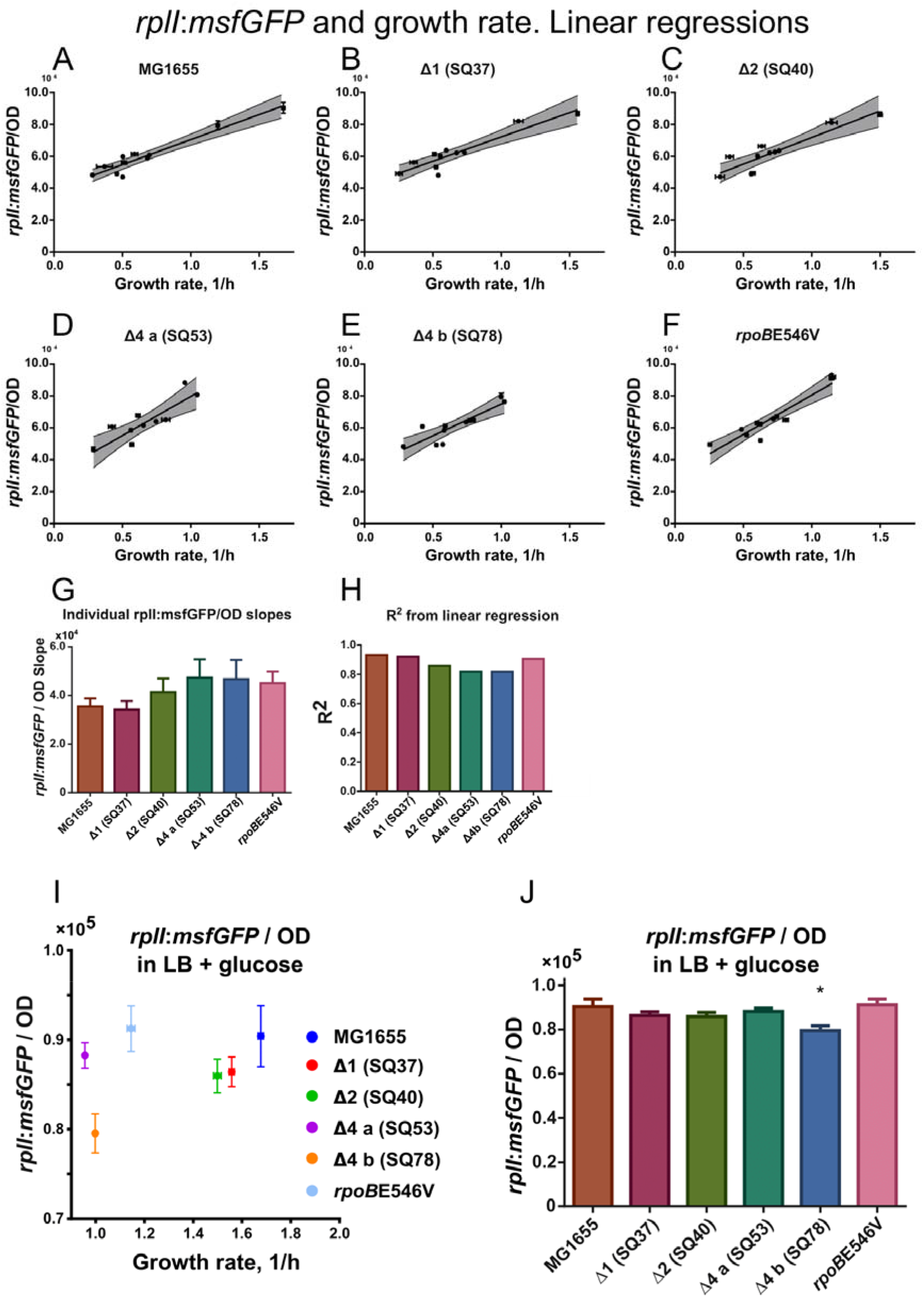
Related to Fig. 2. A-G, Linear regressions of normalized *rplI:msfGFP* as a function of growth rate for mutants grown in different media. Shadowed area represents the 95% confidence interval. H, Bar plot of slopes obtained through linear regression of the rplI:msfGFP/OD values for each growth medium for every strain. I, R2 values representing goodness of fit. I, Normalized fluorescence for strains grown in LB + glucose as a function of growth rate. J, Normalized fluorescence comparison between mutants and the WT in LB + glucose. Error bars represent standard deviations. * denotes significant difference with 95% confidence

**Figure S5.**
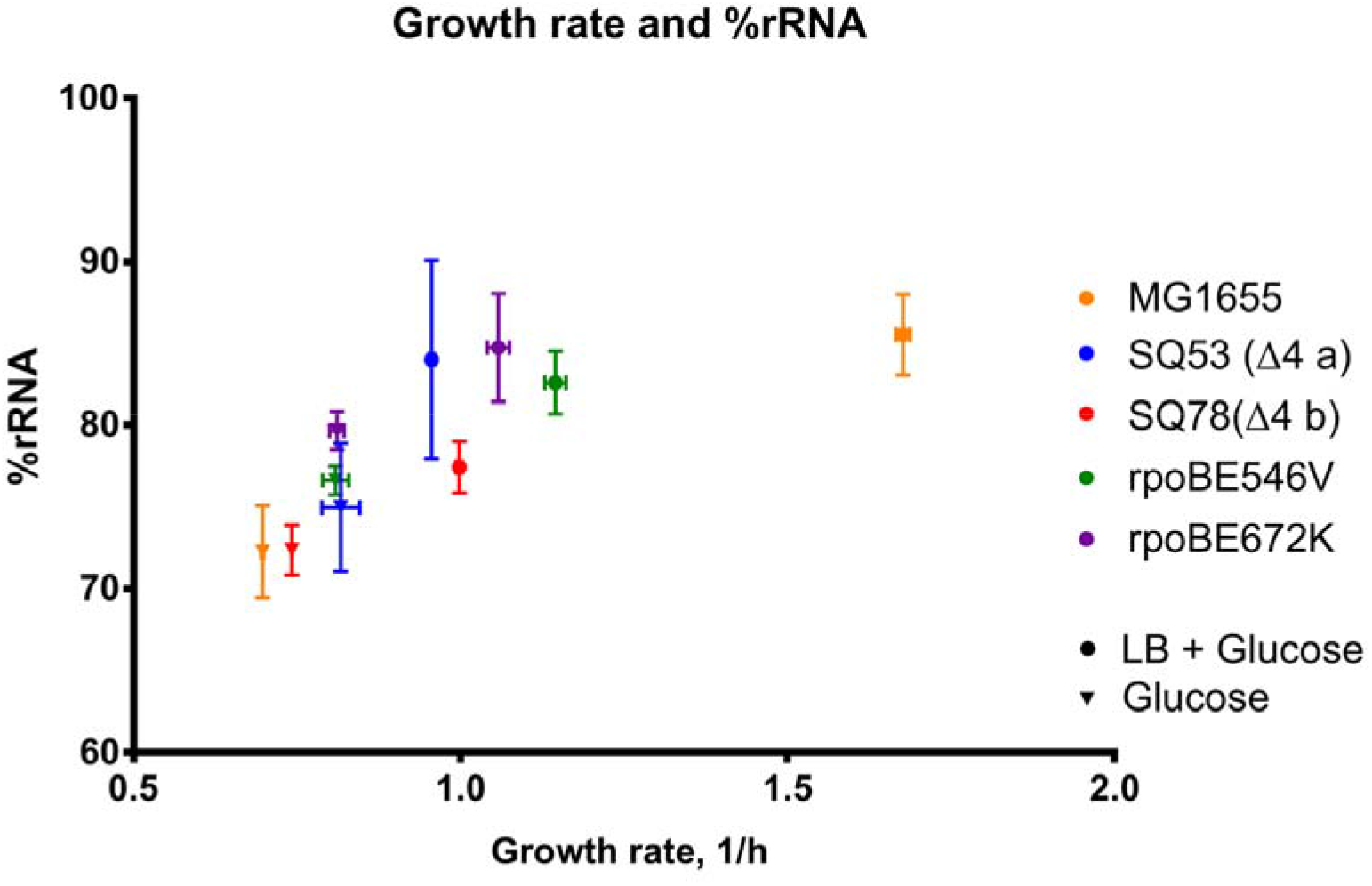
Related to figure 2. %rRNA and growth rates. The % of ribosomal RNA was quantified for selected mutants. rRNA content is increased in mutants in glucose minimal as well as their growth rates. In rich medium, rRNA is similar across strains but mutant growth rates are significantly lower than the WT.

**Figure S6.**
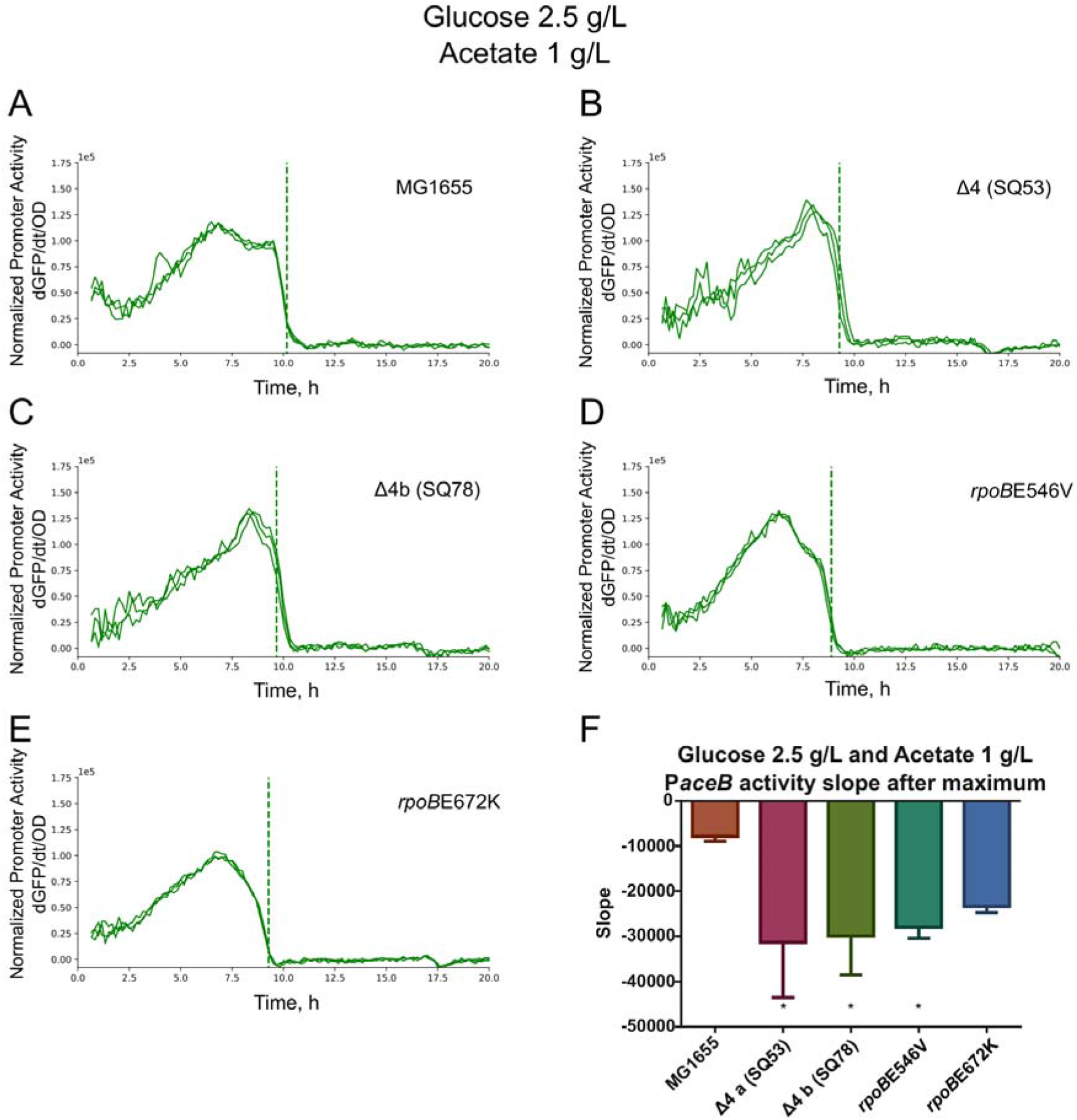
Related to Fig. 3. A-E, PaceB activity in glucose (2.5 g/L) and acetate (1 g/L). Dashed line represents time at growth shifts. F, PaceB activity slopes after maximum. * represents significant differences at 95% confidence.

**Figure S7.**
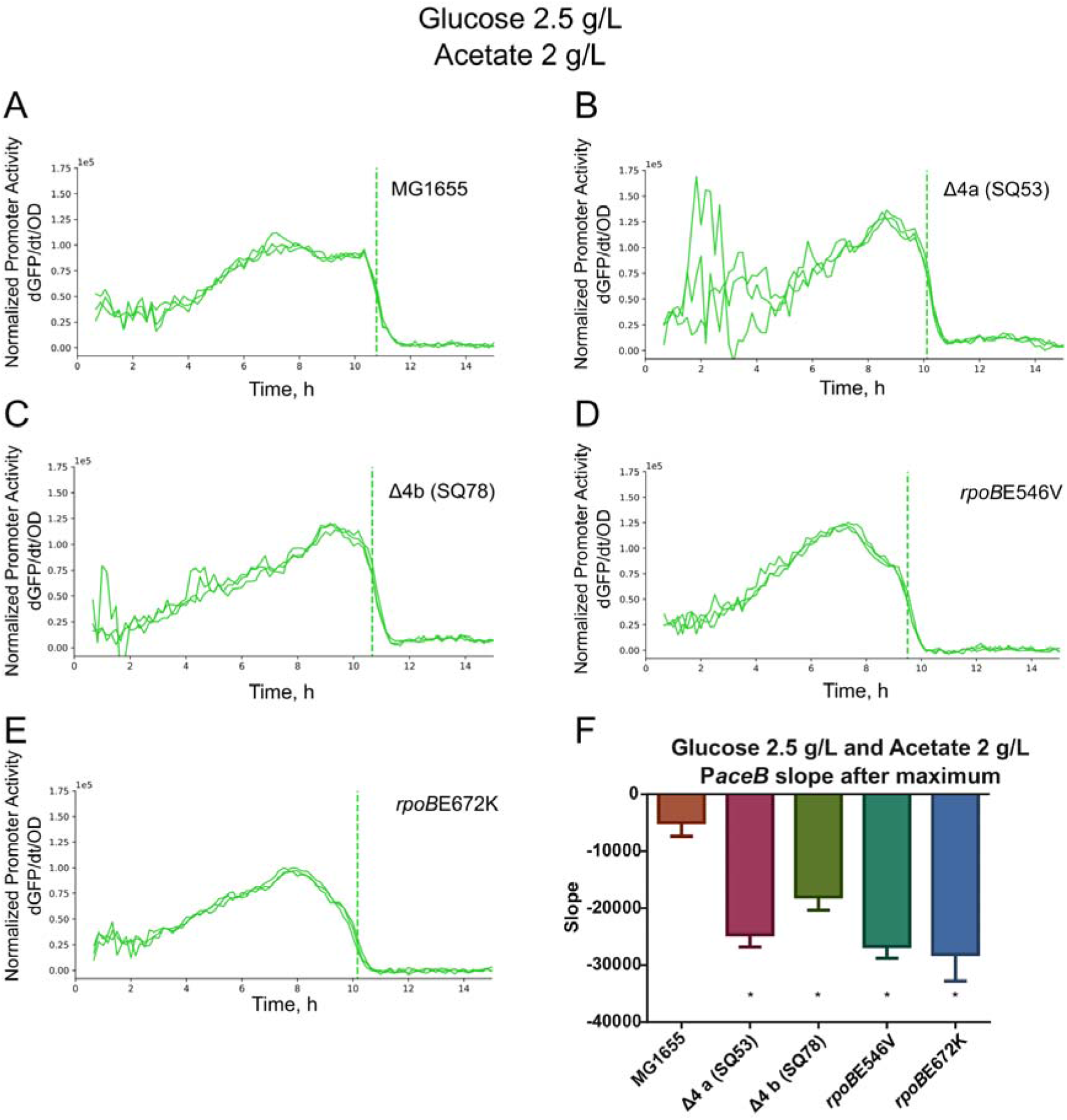
Related to Fig. 3. A-E, PaceB activity in glucose (2.5 g/L) and acetate (2 g/L). Dashed line represents time at growth shifts. F, PaceB activity slopes after maximum. * represents significant differences at 95% confidence.

**Figure S8.**
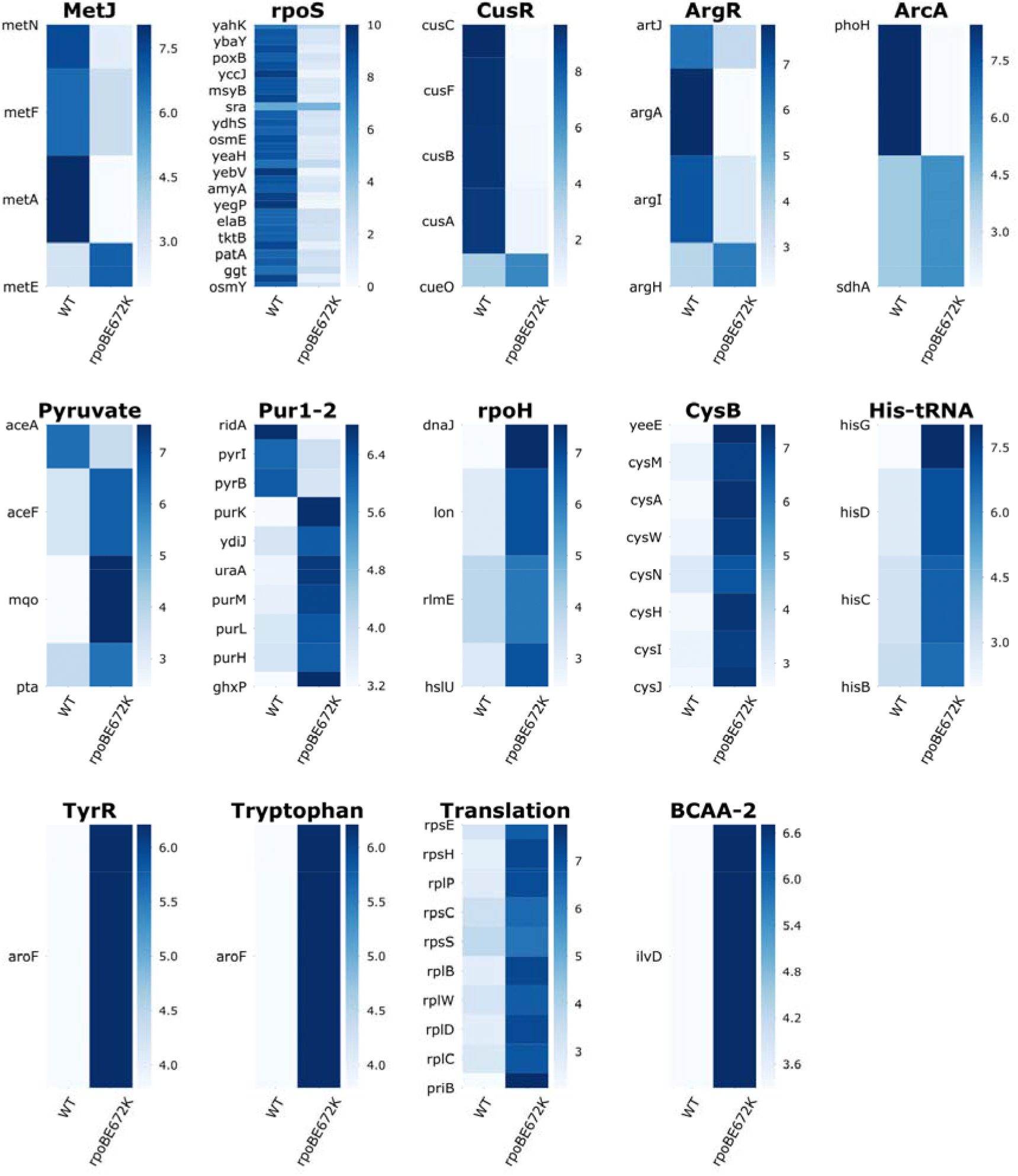
Related to Fig. 6A. The iModulon categorization of rpoBE672K DEGs shows a Fear vs. Greed response, just like SQ53. The number of DEGs in this strain allows for more iModulons to be associated with the transcriptomic data, all of which are shown here. The three iModulons for SQ53 are those presented in figure 6.

